# Bioinformatics analysis and collection of protein post-translational modification sites in human viruses

**DOI:** 10.1101/2020.04.01.019562

**Authors:** Yujia Xiang, Quan Zou, Lilin Zhao

## Abstract

In viruses, post-translational modifications (PTMs) are essential for their life cycle. Recognizing viral PTMs is very important for better understanding the mechanism of viral infections and finding potential drug targets. However, few studies have investigated the roles of viral PTMs in virus-human interactions using comprehensive viral PTM datasets. To fill this gap, firstly, we developed a viral post-translational modification database (VPTMdb) for collecting systematic information of viral PTM data. The VPTMdb contains 912 PTM sites that integrate 414 experimental-confirmed PTM sites with 98 proteins in 45 human viruses manually extracted from 162 publications and 498 PTMs extracted from UniProtKB/Swiss-Prot. Secondly, we investigated the viral PTM sequence motifs, the function of target human proteins, and characteristics of PTM protein domains. The results showed that (i) viral PTMs have the consensus motifs with human proteins in phosphorylation, SUMOylation and N-glycosylation. (ii) The function of human proteins that targeted by viral PTM proteins are related to protein targeting, translation, and localization. (iii) Viral PTMs are more likely to be enriched in protein domains. The findings should make an important contribution to the field of virus-human interaction. Moreover, we created a novel sequence-based classifier named VPTMpre to help users predict viral protein phosphorylation sites. Finally, an online web server was implemented for users to download viral protein PTM data and predict phosphorylation sites of interest.

**Author summary:** Post-translational modifications (PTMs) plays an important role in the regulation of viral proteins; However, due to the limitation of data sets, there has been no detailed investigation of viral protein PTMs characteristics. In this manuscript, we collected experimentally verified viral protein post-translational modification sites and analysed viral PTMs data from a bioinformatics perspective. Besides, we constructed a novel feature-based machine learning model for predicting phosphorylation site. This is the first study to explore the roles of viral protein modification in virus infection using computational methods. The valuable viral protein PTM data resource will provide new insights into virus-host interaction.

## Introduction

Post-translational modifications (PTMs) play a critical role in current proteomics research and regulate protein functions by altering protein interactions, stability, activity, and subcellular localization. Post-translation modifications of viral proteins are relevant throughout various stages of the pathogen life cycle, especially viral infections and genome replication. For example, during entry, the influenza virus carries unanchored ubiquitin chains to engage the host cell’s aggresome system [1]. Once inside the host cell, viral PTMs regulating the infecting process of HSV-1 encode ICP0 protein to degrade host proteins via ubiquitination and sumoylation [2]. In the viral life circle, the HIV-1 Tat protein ser-16 phosphorylated site regulates HIV-1 transcription [3].

Therefore, knowledge of viral PTMs is of great significance to understanding the molecular mechanisms underlying viral infections and recognizing potential drug targets. In recent years, several studies have identified multiple viral PTMs [4-6]; thus, comprehensive analysing these PTM data and establishing a database to provide relevant knowledge is important.

However, few databases have been developed for systematically archiving and easily accessing the PTM sites data of viruses. Also, few researchers have been able to draw on any systematic research into viral PTMs using computational methods. VirPTM [7] stores viral phosphorylation sites and used scan-x to predict modification sites. ViralPhos [8] is a support vector machine based predictor and database that provides outdated viral phosphorylation sites. Bradley et al, studied the phosphorylation motifs in 48 eukaryotes species and 2 prokaryotic species [9]. To date, no databases have collected comprehensive PTM data of viral proteins and few studies analysed the biological significance behind viral PTM data.

To bridge the existing knowledge gap, we have built a viral post-translational modification database (VPTMdb) that first provides comprehensive experimentally verified viral PTM site data, including phosphorylation, sumoylation, glycosylation, acetylation, methylation, ubiquitination, neddylation, and palmitoylation, and it includes 162 studies that have been manually viewed to extract PTM sites. In total, 912 PTM sites from 45 human viruses were obtained, which include 414 manual checked sites from PubMed as well as 498 sites from UniProtKB/Swiss-Prot.

Secondly, by using computational methods, we investigated the PTM sequence motifs, the function of target human proteins, and characteristics of PTM protein domains. This work will generate fresh insight into viral infection mechanisms as well as identify virus PTM sites.

Finally, PTM was predicted in other species with machine learning approaches [10, 11]. For viral protein serine modification site identification, we implemented a novel feature-based classifier named VPTMpre into the VPTMdb to provide users with the ability to find viral protein phosphorylation sites. The results of independent testing showed that VPTMpre represents a powerful tool to predict viral protein phosphorylation sites.

The online web server is available at http://vptmdb.com:8787/VPTMdb/, and users can browse and download viral PTM data freely. Support vector machine, random forest, and naïve Bayes were integrated into VPTMpre, and users are able to choose one machine learning model to predict possible phosphorylation sites of interest.

## Results

### Database contents

**Fig 1** shows that the VPTMdb web server consists of two parts: VPTM database and VPTMpre. The VPTM database currently includes 414 unique experimentally determined PTM sites with 8 modification types from 45 viruses. In summary, 162 manually checked references were collected in the database. Each entry in VPTMdb includes the (i) virus name, (ii) virus protein name in the UniProt database, (iii) PTM type, (iv) viral modification site, (v) residue sequences, (vi) kinase, (vii) a short description of the PTM site extracted from the publication, and (viii) PubMed id. PTM data from UniProtKB/Swiss-Prot contain two types: 199 phosphorylation sites and 299 glycosylation sites (N-lined and O-lined).

**Fig 1.**
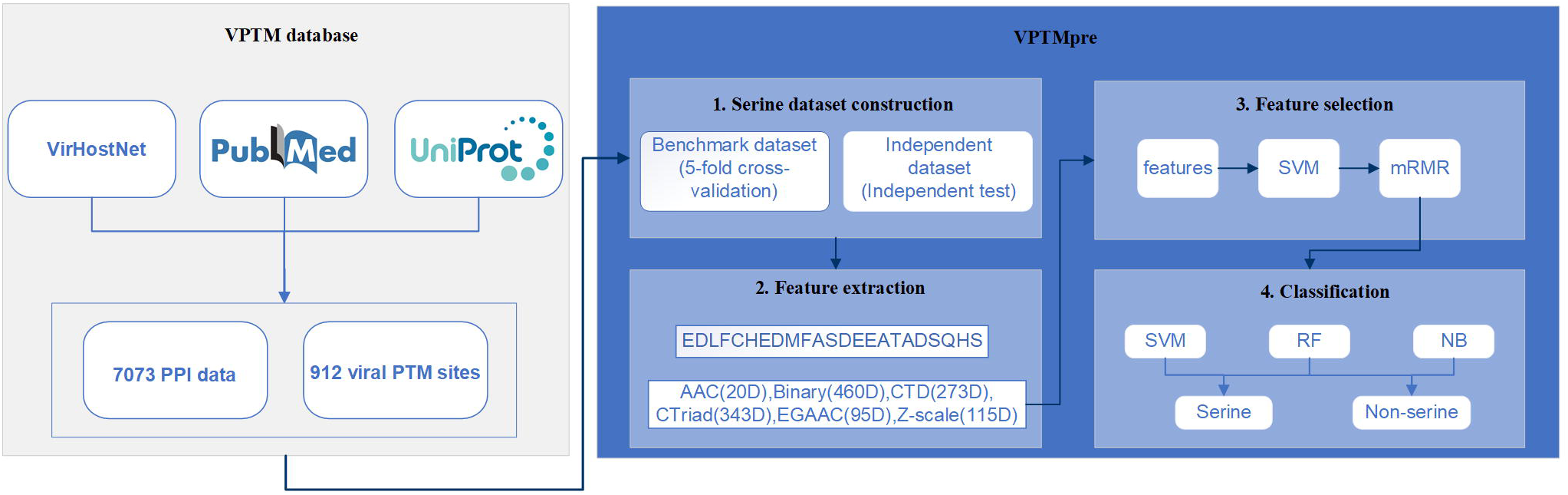
Overview of VPTMdb. Framework of VPTMdb web server construction. First, PTM data were collected from PubMed and UniProt/Swiss-Prot. Then, VPTMpre was constructed to predict viral protein phosphorylation sites.

The statistics of experimentally verified sites in VPTMdb show that among eight PTM types, phosphorylation sites account for the most (484 sites, including 285 manually checked and 199 sites from UniProtKB/Swiss-Prot) at more than 50% of the total database. The top five viruses in the number of manually checked modification sites are HAdV-2 (51 phosphorylation sites), EBOV (29 phosphorylation, 1 sumoylation, 2 ubiquitination, 8 acetylation sites), HIV-1 (21 phosphorylation, 4 sumoylation, 2 ubiquitination, 5 acetylation and 3 glycosylation sites), H1N1 (19 phosphorylation, 3 sumoylation, 2 ubiquitination, 6 acetylation and 2 glycosylation sites), and HCV (10 phosphorylation, 1 sumoylation, 1 ubiquitination, 1 methylation 4 palmitoylation and 14 glycosylation sites) (**S1 Fig**).

Human-virus PPI data were included in the VPTMdb, which are helpful to determine the potential function of PTMs during viral infections. PPI data in the VPTMdb contains 7073 interactions with 2934 proteins in 43 viruses. **Fig 2** shows the distribution of modified proteins in the protein-protein interaction network.

**Fig 2.**
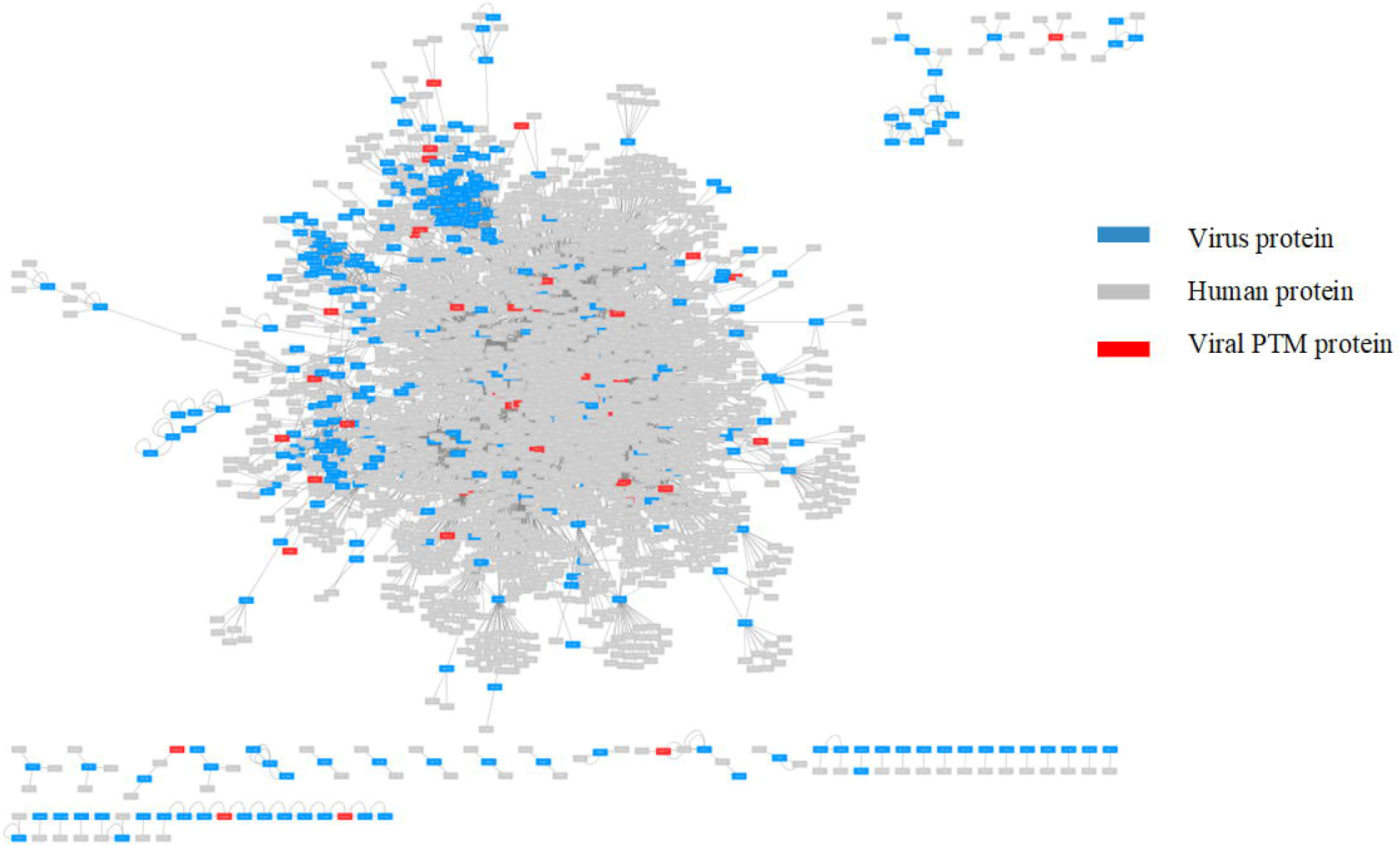
The virus-human protein-protein interaction network. Each node represents viral protein or human protein. Each edge represents virus-human or virus-virus association.

The web server involves five easy-to-use main pages: ‘Home’, ‘Browse’, ‘Prediction’, ‘Download’, and ‘Help’. Each of these pages enables users to search, browse, predict, and download data without any prerequisite knowledge. In the ‘Browse’ section, users can search the PTM data conveniently by typing keywords in the search box and download data freely, what is more, virus-human protein-protein interaction data are provided and visualized. The ‘Prediction’ page provides VPTMpre, a sequence-based machine learning predictor for phosphorylation serine site prediction. All data about virus PTM are stored in the ‘Download’ page for batched downloading. The ‘Help’ page contains a detailed tutorial to help users learn about VPTMdb.

### Investigation of viral PTM sequence motifs

Previous research has reported that most eukaryotic species have universal kinase-substrate motifs in their phosphorylation proteins [9]. The human viruses are living in the cell, and their proteins are modified by human kinase or viral protein kinase. To this end, we were interested in a question: Are the modified substrate motifs of viral proteins the same as human proteins motifs? To answer this question, we used the motif-x tool [12] to extract motifs from viruses.

As shown in **Fig 3**, for viral phosphorylation modified proteins, when kinases were from human proteins, the viral sequences motifs were the same as human proteins (xSPx) (“x” means any residue) [9]. For viral protein SUMOylation, we noted that the highly prevalent motif across 16 viruses was KxE, which was also enriched in human proteins [13]. What’s more, we investigated viral N-glycosylated proteins’ motifs. The results showed that NxS/T is the significant motif.

**Fig 3.**
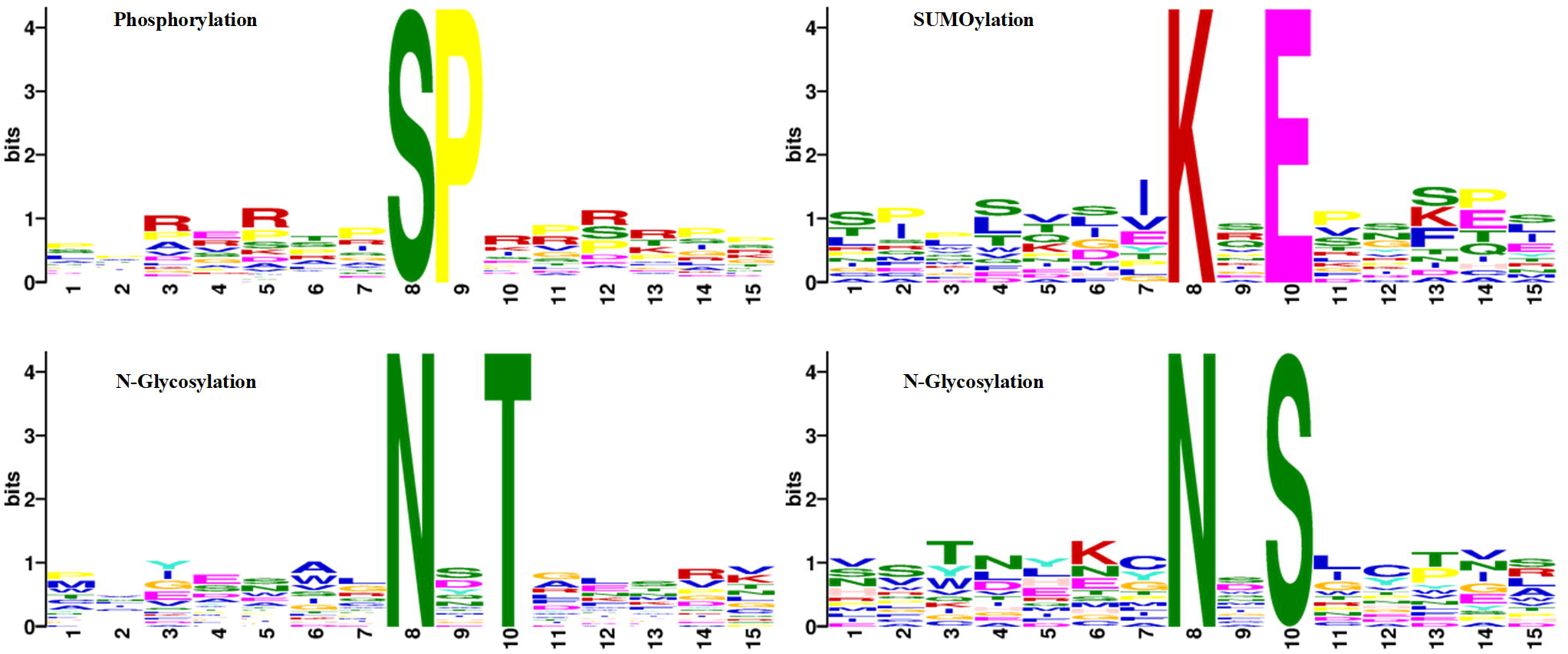
Viral protein PTM motifs discovered by motif-x.

We also investigated protein motifs when kinases were viral proteins. In VPTMdb, 13 amino acid residues were modified by viral protein kinases (HSV-1 US3 or HSV-2 UL13). However, there are no significant motifs when used motif-x tool. Thus, sequence logo was used to visualize PTM sequences (**S2 Fig**). Unlike human protein kinases, arginine (R) was enriched near the serine site modified by virus kinase.

Overall, these results suggest that the phosphorylation, SUMOylation, and N-glycosylation residues in viral PTM sequences have the consensus sequence motifs with human PTM proteins. Viruses may use those short motifs to interact with human proteins and utilize human signal pathways to regulate themselves replication.

### Function characterization of viral PTM protein target human protein

To investigate how viral PTM proteins influent the human cellular activities, we created virus-human protein-protein interactions (PPI) network. The virus-human PPI data consist of virus-human and virus-virus interactions (viruses are these in VPTMdb database). PPI network includes 2934 proteins and 7073 interactions. The degree was considered as the metric to evaluate the role of viral proteins in the virus-host PPI network.

Firstly, the roles of viral PTM proteins in the PPI network were analysed. Notably, in Influenza A virus(H1N1), HPV-18, HPV-31, HPV-8, HIV-1, HTLV-1, EBOV, SARS-Cov, hRSV, and Vaccinia virus, their all PTM proteins have significant large degrees than average network degrees (**S1 Table**).

Then, the Gene Ontology and KEGG enrichment analysis were performed to characterize the function of target human proteins, which may reflect how viral PTM proteins influent human cellular activities. It is interesting to see that the top five enriched KEGG pathways were “Ribosome”, “Spliceosome”, “Proteasome”, “RNA transport” and “Mismatch repair”. It reveals that viruses use human proteins to promote their transcription and modifications. Also, it has been observed that the top ten GO enrichment terms were related to protein targeting, translation, and localization (**S3 Fig**).

### Viral PTMs are more likely to be enriched in protein domains

We analysed the domain composition of viral PTM protein. The protein domain data were extracted by HMMER, then 141 domains were obtained and 62 out of 141 domains have modified residues. These domains which have PTM sites were from 57 proteins in 30 viruses. We counted the number of modifications on proteins in the 30 viruses and found that 53.4% of the modifications were distributed in PFAM protein domains. On average, there are 1.33 modification sites per 100 amino acids for the viral PTM proteins, which increased to 2.1 modification sites per 100 amino acids for the viral PTM domains. These results indicated that viral PTMs are more probably enriched in protein domain regions.

### Feature-based predictor construction

For viral protein phosphorylation site prediction, we used the feature representative strategy to create a novel classifier. The first step is to compare different features and evaluate their predictive power. The data in **Table 1** show that six features as well as their combinations were evaluated in SVM with a 5-fold cross-validation. AUC, F1-score and MCC were used as the performance evaluation indicators. The results declare that the z-scale, which captures the physical-chemical information of amino acids, is the best among the six single features (AUC=0.957, F1-Score=0.887, MCC=0.810). For BINARY, EGAAC and CTriad, their AUC values also achieved above 90.00%. Moreover, when we fused the features, the result showed that ZSCALE combined with AAC features improved the sensitivity, F1-score and AUC by 8.40%, 1.5%, 0.1% compared with individual z-scale features.

**Table 1.** Comparison of performance between the single features and fused features with the mRMR method.

| Features | Dim | Sn | Sp | MCC | F1 | AUC |
| --- | --- | --- | --- | --- | --- | --- |
| **1.AAC** | 20 | 0.738 | 0.739 | 0.479 | 0.738 | 0.821 |
| **2.BINARY** | 460 | 0.827 | 0.896 | 0.732 | 0.857 | 0.931 |
| **3.ZSCALE** | **115** | **0.812** | **0.985** | **0.810** | **0.887** | **0.957** |
| **4.EGAAC** | 95 | 0.896 | 0.850 | 0.747 | 0.876 | 0.901 |
| **5.CTDD** | 195 | 0.996 | 0.077 | 0.184 | 0.682 | 0.655 |
| **6.CTDC** | 39 | 0.735 | 0.696 | 0.433 | 0.720 | 0.795 |
| **7.CTDT** | 39 | 0.823 | 0.712 | 0.541 | 0.779 | 0.827 |
| **8.CTriad** | 343 | 0.823 | 0.870 | 0.694 | 0.843 | 0.926 |
| **{1,3}** | **135** | **0.896** | **0.908** | **0.806** | **0.902** | **0.958** |
| **{2,3}** | 575 | 0.873 | 0.888 | 0.764 | 0.881 | 0.94 |
| **{3,4}** | 458 | 0.835 | 0.904 | 0.743 | 0.866 | 0.944 |
| **{3,8}** | 210 | 0.873 | 0.896 | 0.771 | 0.885 | 0.943 |
| **{3,5,6,7}** | 388 | 0.831 | 0.85 | 0.681 | 0.839 | 0.921 |
| **{2,3,4,8}** | 1013 | 0.85 | 0.908 | 0.762 | 0.876 | 0.947 |
| **{1,2,3,8}** | 938 | 0.742 | 0.985 | 0.751 | 0.844 | 0.938 |

Note: The first column represents the different feature extraction methods employed in this study. Dim refers to the different dimensions of every feature, and Sn, Sp, MCC, F1 and AUC represent the sensitivity, specificity, Mathews Correlation Coefficient and AUC value, respectively.

However, the combination of EGAAC, BINARY, ZSCALE and CTriad features did not significantly enhance the model’s performance, which suggests that high-dimensional features may include useless features that weaken the model performance. Among all the features, considering the three evaluation values of F1-score, MCC, AUC and dimensions, the AAC combined with the ZSCALE performed best, and the sensitivity, AUC and F1-score were higher than the single z-scale features. The independent test also shows that AAC combined with ZSCALE features significantly increased the AUC, F1-score, MCC, and Sn by 0.90%, 21.7%, 2.60%, and 25.0%, respectively (**S1 Supporting Information**).

Now, it is important to answer two questions: (i) what is the difference between phosphorylation sites and non-phosphorylation sites and (ii) which features contribute most to the viral phosphorylation protein? To this end, we analysed the z-scale feature information between phosphorylation sites and non-phosphorylation sites. Then, we selected the most important features from the combined features with the mRMR method and using svm, random forest and naïve Bayes to perform a predictive evaluation.

### Z-scale feature analysis

The z-scale feature based on amino-acids’ physical-chemical properties includes five z values. The distribution of amino acid residues around serine sites is able to determine the different physicochemical properties between phosphorylation sites and non-phosphorylation sites. From **Fig 4**, we can see that the z3 values of the phosphorylation sites are smaller than that of the non-phosphorylation sites, implying that a more negative charge occurred around viral protein phosphorylation sites than around non-phosphorylation sites. The results also showed that the z1, z2, z4, and z5 values of the phosphorylation sites are bigger than that of the non-phosphorylation sites. Overall, the different z-scale compositions surrounding the phosphorylated and non-phosphorylation sites indicate that it is reasonable to choose the z-scale as a feature for prediction.

**Fig 4.**
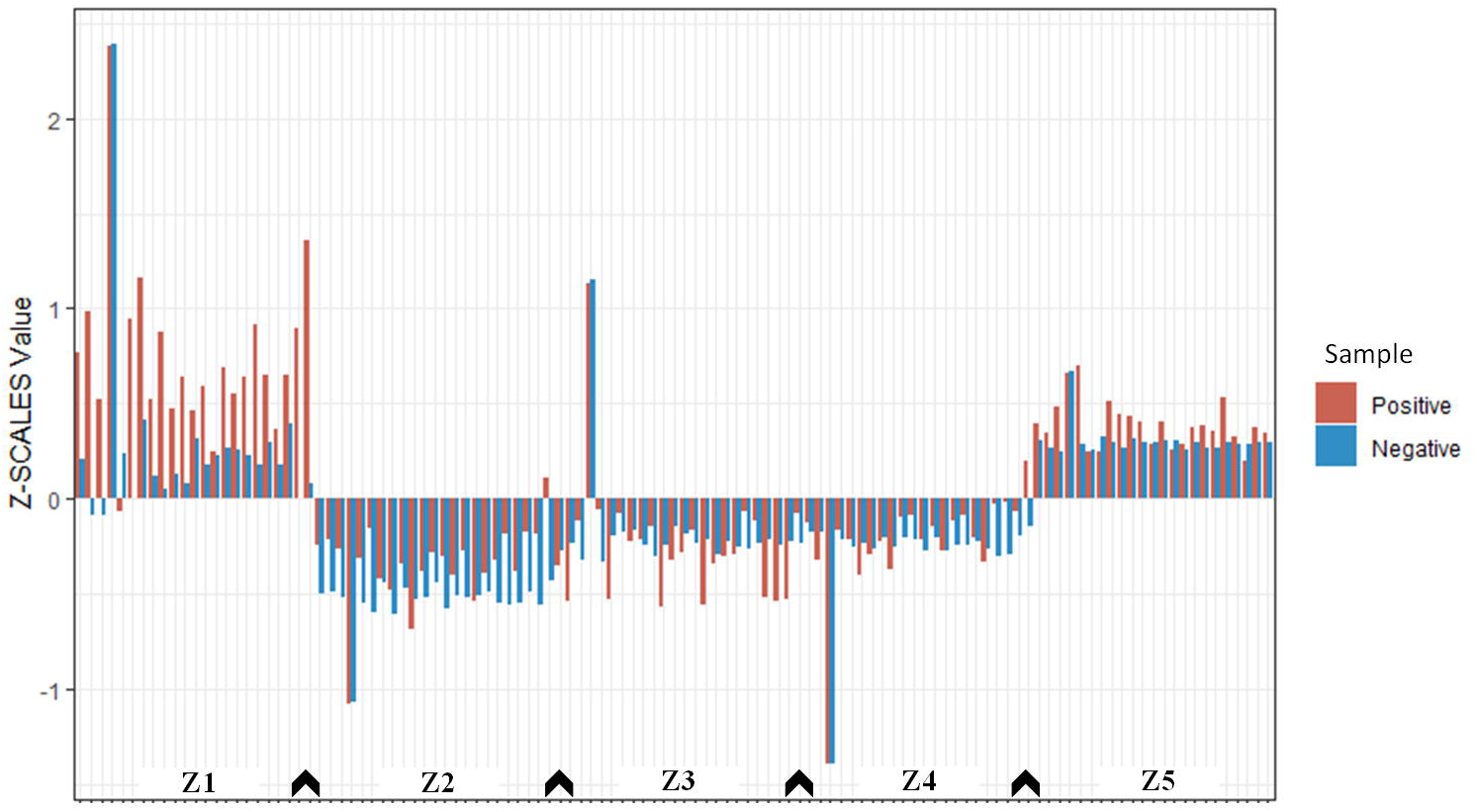
Comparison of the z-scale in positive and negative datasets. The vertical axis represents the z-scale values. The X-axis represents the five binary sequences.

### Performance evaluation

VPTMdb provides three classifiers: support vector machine, random forest and naïve Bayes. Different dimensional features may have different impacts on different predictors. Thus, we selected features of different dimensions using the mRMR algorithm and compared the three classifiers’ performance from the 5-fold cross validation (**S1 Supporting Information**).

**Fig 5A** shows that the maximum AUCs of the svm and random forest are similar. For the random forest and svm, the AUCs increased when more features were selected (random forest: 14-135 features, with AUC > 0.90; svm: 27-135 features, with AUC > 0.90). However, we observed that the AUCs of naïve Bayes (AUCs > 0.80) decreased when more features were added. From a statistical point of view, to prevent the curse of dimensionality, fewer and more meaningful features should be chosen. Taking the above results into consideration, for 68 features, the AUCs of the three predictors perform better, suggesting that 68D is the most meaningful feature among all the features.

**Fig 5.**
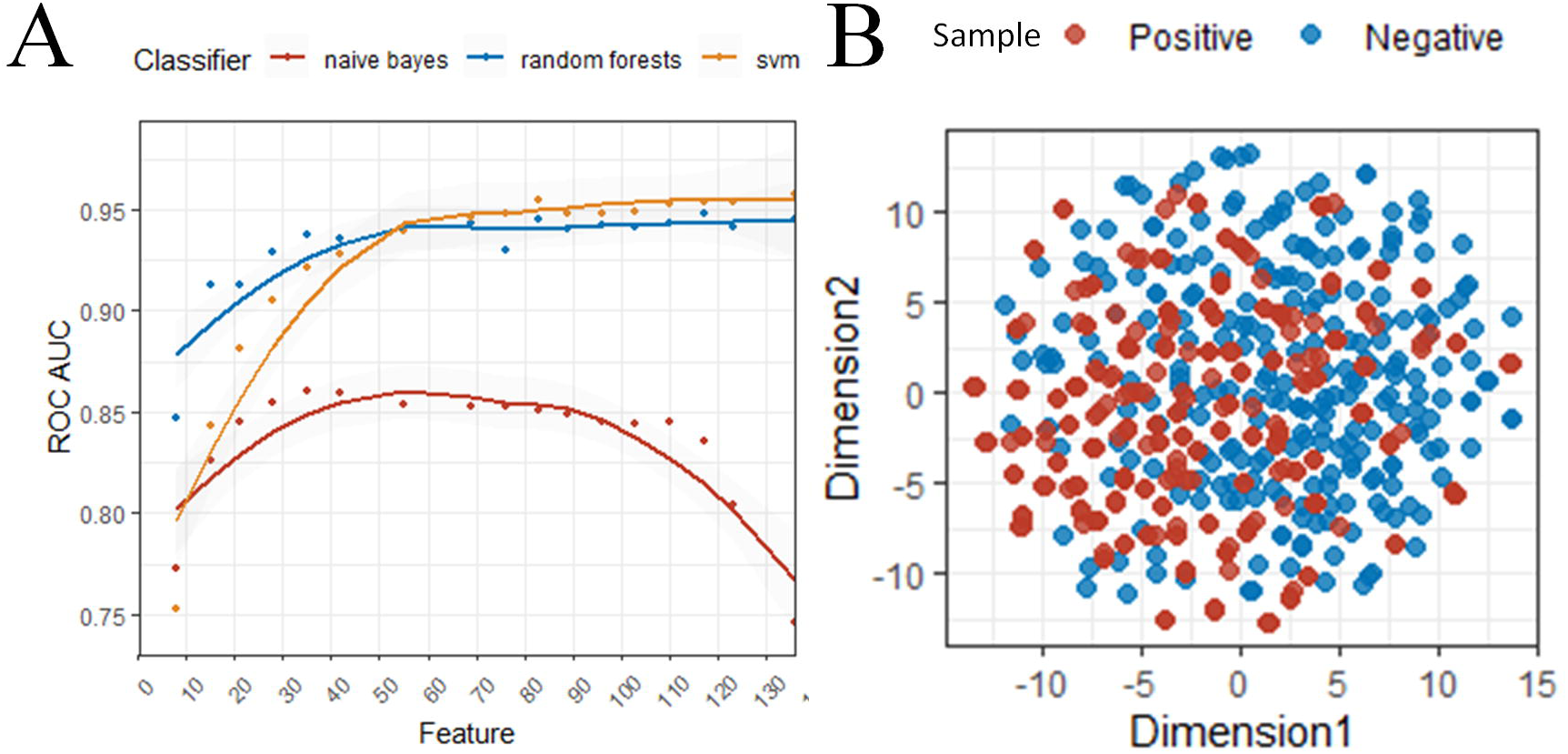
Feature-based predictor construction. (A) Five-fold cross-validation performance of the three classifiers on different features. (B) t-SNE visualization of positive and negative data using 68D features.

To understand the effective of our 68-dimensional features, the T-distributed Stochastic Neighbour Embedding (t-SNE) algorithm was used to visualize the positive and negative samples. A clear distinction was observed between the positive and negative samples, implying that our features selection results are effective (**Fig 5B**).

To assess the robustness and performance of the svm, random forest, and naïve Bayes in 68D features, 10-fold random independent tests were performed. The model performance on independent datasets is shown in **Fig 6**, random forest performed better, the average AUC, MCC, F1-score of its are 0.744, 0.427, 0.656 respectively. Comparing random forest and PSI-blast (**S1 Supporting Information**), the MCC, acc and sp values of random forest are higher than PSI-blast for 6.92%, 2.8% and 19.1%. Taking all indicators into consideration, our method is stable and better performance. We implemented svm, random forest and naïve Bayes into VPTMpre, users can choose them to predict phosphorylation sites of interest.

**Fig 6.**
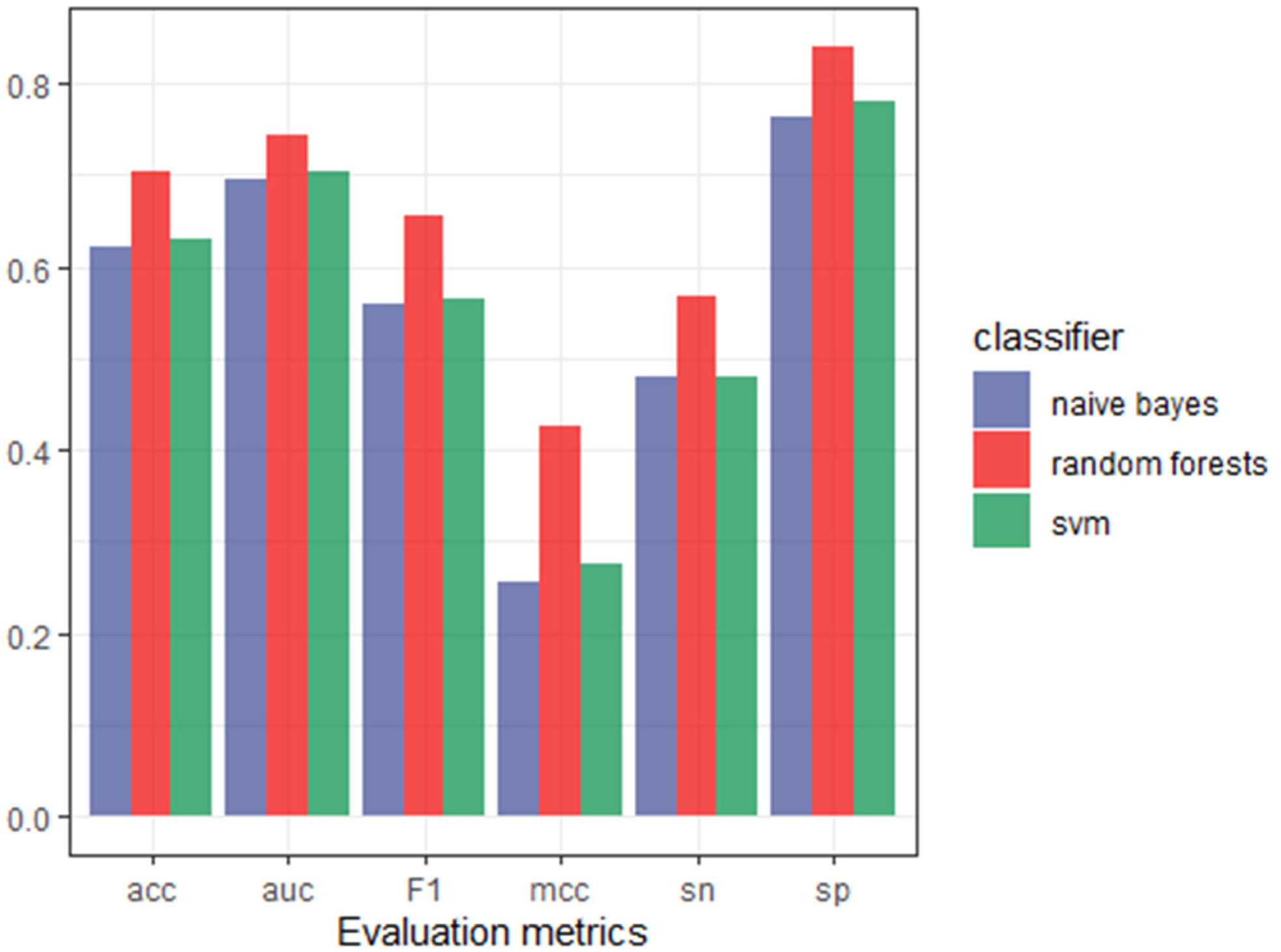
Independent test results. Sensitivity, specificity, AUC, MCC and F1-score of the proposed features in three classifiers.

## Discussion

In this work, we constructed VPTMdb, which is the first database that systematically collected experimentally verified viral protein PTMs. Virus-human PPI data were also collected in the VPTMdb to determine PTM sites association functions. These viral protein PTM data provide unique insights into virus-host interactions.

Firstly, viruses in VPTMdb have the same substrate motifs as human proteins in phosphorylation (37 viruses), SUMOylation (16 viruses) and N-glycosylation (6 viruses). Several studies have shown that viral functional motifs play significant roles in virus life cycles and virus-host interactions [14]; For instance, SUMOylation motifs can promote viral proteins binding and enhance viruses replication as well as immune evasion [15, 16]. Hence, these conserved sequence motifs in viral proteins may help them to hijack host PTM processes and utilize cellular substance to facilitate virus infections.

Secondly, the function of the viral PTM proteins target human proteins were explored. The results showed that ten viruses PTM proteins have more degrees than the network average degrees. One possible reason is that viral proteins modification processes require the cooperation of multiple other proteins, so modified proteins have more interaction partners. Another possible reason is that PTMs regulate the state of proteins, and modified proteins can perform more functions. For instance, HCV core protein represses transcription of p21 is regulated by the phosphorylation at serine-116 site [17]. These PTMs will significantly change the function and interaction partners of viral proteins. Also, the top ten GO enrichment results of target human proteins were related to binding, which was partially validated that PTM proteins tend to bind with more human proteins.

Moreover, we found that viral PTM sites are more likely to be enriched in the protein domains; Studies have shown that human modified lysines are more likely near phosphorylation sites, which form a PTM cluster region [18]. For viruses, these cluster PTMs in protein domains may form short motifs to enhance the regulate function of viral proteins.

Finally, based on the analysis of viral PTM protein features, VPTMpre, a novel feature representative classifier, was developed to predict viral protein serine sites. We compared various feature extraction methods and selected the optimized features using the mRMR algorithm. The feature analysis results showed that 68D was able to distinguish the phosphorylation sites and non-phosphorylation sites in viral proteins. VPTMpre was integrated into the VPTMdb web server to provide an online phosphorylation site prediction service. Users can choose three classifiers (svm, random forest and naïve Bayes) to predict phosphorylation sites of interests. However, because of data limitations, the prediction of VPTMpre is limited to serine sites. With a continuous collection of new viral PTM data, we expect that VPTMpre will be extended to predict more types of PTM sites and obtain a better performance.

In the future, to respond to the rapid growth of viral PTM data, VPTMdb will be updated regularly and more viral PTM-related data collected to ensure that it provides the most comprehensive information to users. As the first attempt to develop the comprehensive viral PTM database, we sincerely welcome support and suggestions from the research community to improve the VPTMdb database.

## Methods

### Data collection

There are three major steps in data collecting and pre-processing, which are described below.

Firstly, we queried PubMed using the keyword search terms: (virus name) and (eight modification types) for studies published before Jan 01, 2020. As a result, 6052 papers were obtained, each of which was manually retrieved using the following standards: (i) the viral post-translational modifications were experimentally verified; and (ii) if two references contained the same PTM site, the earliest published study was retained. In total, 45 viruses, 162 papers and 414 PTMs were obtained.

Subsequently, 498 viral PTM data points from UniProtKB/Swiss-Prot were integrated into VPTMdb. For experimentally validated virus PTM types, the sites were extracted manually from the articles mentioned above. The protein sequences, UniProt ID and PMID were mainly extracted from NCBI, UniProt and PubMed. Finally, human-virus protein-protein interactions were collected from the VirHostNet based on viral strains in the VPTMdb.

### PTM data analysis

The phosphorylation (37 viruses), SUMOylation (16 viruses) and N-Glycosylation (6 viruses) data were from VPTMdb. Motif-x tool was employed to extract motifs using its default parameters (score-threshold of 1 × 10^−6^, min-occurrences of 5, and width of 15). Proteins domains were searched by HMMER (using PFAM database) with default parameters. PPI data were downloaded from VirHostNet database. Gene Ontology and KEGG enrichment analysis used clusterProfiler [19]. Network analysis was performed using Cytoscape [20].

### Overview of viral phosphorylation sites prediction

Identifying viral protein PTM sites by experimental methods is still expensive and time consuming. Thus, predicting them in *silico* using bioinformatics approaches is necessary. To this end, a sequence-based classifier named VPTMpre was created to predict viral post-translational modification serine sites. Because threonine and tyrosine data are too few to train the model, we only predicted serine sites in this study.

Five main procedures were performed to build the VPTMpre predictor. (i) a balanced benchmark dataset was constructed using the Synthetic Minority Oversampling Technique (SMOTE) [21] sampling method (**S1 Supporting Information**); (ii) various feature representative methods were compared to obtain an effective feature representation strategy, with support vector machine used as the base classifier in a 5-fold cross-validation approach to find the best feature groups; (iii) the predictive performance of three classifiers (svm, random forest, naïve Bayes) on different feature dimensions was compared using the Minimum redundancy and maximum relevance (mRMR) method, and the features that performed well in all three classifiers were selected as the most meaningful and significant features; (iv) a 10-fold random independent test was performed to evaluate the predictive performance of the three different classifiers (svm, random forest, naïve Bayes); and (v) VPTMpre was implemented in the online web server.

### Data preparation and processing

All viral phosphorylation experimentally verified serine sites in our database were used as positive samples, and those not marked by any phosphorylation information on the same protein were considered negative samples. As a result, we obtained 182 phosphorylated serine residues as well as 2148 non-phosphorylated residues. Phosphorylation sites from UniProtKB/Swiss-Prot were regarded as the independent dataset, and they included 93 positive serine sites and 1878 negative serine sites. After using CD-HIT (clustering thresholds set to 0.8) [22] to remove redundant sequences, we obtained 129 positive sites and 1611 negative sites. The independent dataset contained 52 positive sites and 1072 negative sites (**Table 2**). These sequences were truncated to a 23-residue symmetrical window (−11 to 11).

**Table 2.** Summary of training and independent datasets.

| Datasets | Types | Total number | After deletion | After balanced |
| --- | --- | --- | --- | --- |
| Training set | Positive | 182 | 129 | 260 |
|  | Negative | 2148 | 1611 | 260 |
| Independent set | Positive | 93 | 52 | 52 |
|  | Negative | 1878 | 1072 | 1072 |

In order to eliminate the prediction bias caused by data imbalance, we re-sampled the training data by SMOTE methods and obtained 260 positive sites and 260 negative sites, which consisted of the training dataset. The negative test set from UniProtKB/Swiss-Prot was randomly divided into twenty parts (**S2 Table**). We randomly select ten negative subsets from the twenty parts and combined them with ten replicate positive sets to constitute ten independent test datasets (**S1 Supporting Information**).

### Feature representation

To achieve a better classification effect, a key step is feature extraction, which means that a protein sequence is encoded as a numeric vector for machine learning model.

#### Amino acid composition (AAC)

AAC is the frequency of 20 amino acids for a given sequence [23]. This descriptor can be denoted as follows:

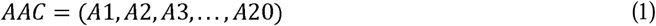

Where

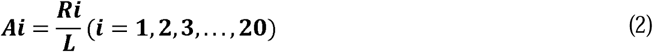

*Ri* is the observed number of types i amino acid in a protein sequence. L is the length of protein. Thus 20 features were obtained, and sum of which is 1.

#### Binary profile

The binary profile transformed each amino acid into a 20-dimensional binary numerical vector. For instance, the alanine (‘A’) is deciphered as 10000000000000000000, cysteine (‘C’) is deciphered as 01000000000000000000, etc. Consequently, we obtained a 460-dimensional vector for this binary profile feature.

#### Conjoint triad (CTriad)

The conjoint triad feature is sequence information for proteins. Twenty amino acid types are clustered into seven classes to construct the C-triad feature.

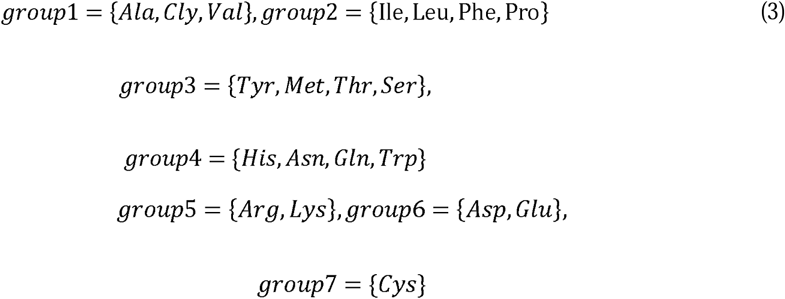

First, protein sequences are encoded into a numerical vector using the AA groups list above. Subsequently, any three continuous AAs are regarded as a unit, and scanning along the sequences and counting the frequencies of each triad type is performed to obtain a 343-dimensional numerical vector.

For example, a protein sequence S contains L AA residues, which are expressed as follows:

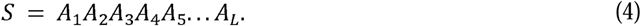

Then, we scan along the sequence with a slide window in three continuous residues:

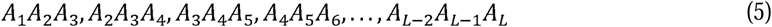

Finally, the C-triad feature of a protein is defined as the frequency of the corresponding triad type in that protein:

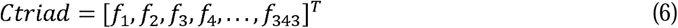

where,

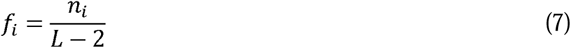

*n*_i_ is the occurrence number of the i-th triad type (i= 1, 2, …, 343). More detailed information about C-triad can be found in [24].

#### Composition-Transition-Distribution (CTD)

CTD clusters 20 amino acids into three groups: hydrophobic, neutral and polar. The CTD composition (CTD-C) calculates the composition values of hydrophobic, neutral and polar groups for a given sequence. The CTD transition (CTD-T) represents the percentage frequency of an amino acid of one particular property followed by an amino acid of another property. The CTD distribution (CTD-D) represents the distribution of each property for a given sequence. Each property has five distribution descriptors, which are the first residue, 25% residues, 50% residues, 75% residues, and 100% residues in the whole sequence of a given specific property. In this research, CTD-C, CTD-T, and CTD-D were used to encoded protein sequences and yielded 39, 39, and 195 features, respectively. More detailed information about CTD can be found in the literature [25].

#### Enhanced grouped amino acid composition (EGAAC)

EGAAC was first proposed by Chen et al. [26] and is the improved version of GAAC features. GAAC divides 20 standard amino acids into five groups based on their physical and chemical properties. The formulation of GAAC is as follows:

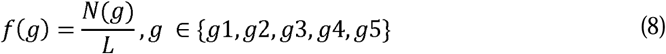

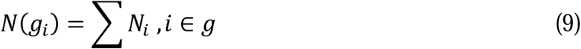

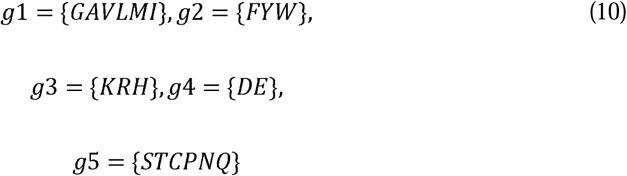

where *L* is the length of sequence, *N(g)* is the number of amino acids in group g, and *N*_*i*_ is the occurrence number of i-th amino acid type.

EGAAC scans along the sequence and calculates the GAAC values in a fixed-size window:

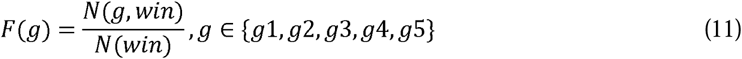

where *N(g,win)* is the number of amino acids in group g within a fixed-size window *win* and *N(win)* is the window size. *win* ranges from 1 to 17. In this study, the window size was set to 5, and we finally obtained a 95-dimensional vector.

#### Z-Scale (ZSCALE)

Z-scale is a feature descriptor that describes AAs’ physicochemical properties. It was first published by Hellberg [27], who introduced three z-scales (z1-z3), and then Sandberg et al. (Sandberg, et al., 1998) improved the original z-scale features by adding two more z-scale values, using 26 properties of 87 AAs. In this study, we employed the z-scale using five scales(z1-z5). The five z-scales are based on lipophilicity (z1), bulk (z2), polarity/charge (z3), electronegativity and heat of formation(z4), electrophilicity and hardness(z5), yielding a 115-dimensional numerical vector.

### Feature selection and optimization

Generally, high-dimension biological features may be noisy, which led to poor prediction performance. However, feature selection is a good strategy to overcome feature redundancy. Feature selection means using a reduction algorithm to select the major features that are able to improve the performance of specific classifiers.

In this work, six descriptors and their combined features’ performance were compared using 5-fold cross validation in the training data with the Support Vector Machine (SVM) method. Subsequently, the Minimum redundancy and maximum relevance (mRMR) method was chosen to select the most meaningful features. To investigate the predictive performance of three classifiers, we compared the different dimensions of features in the svm, random forest, naïve Bayes methods. The features that performed well in all three classifiers were selected as the most meaningful and significant features. The T-distributed Stochastic Neighbour Embedding algorithm was used to visualize the features[28].

### Performance evaluation

Sensitivity (Sn), Specificity (Sp), F1-score, and Mathews Correlation Coefficient (MCC) were applied to estimate the prediction performance (**S1 Supporting Information**). Besides, the receiver operating characteristic (ROC) curve and the area under the ROC curve (AUC) were used to evaluate the overall performance of the model. The ROC curve is a continuous line plotted by the false positive rate (FPR) as the X-coordinate and true positive rate (TPR) as the Y-coordinate. The higher the AUC value, the better the performance of the classifier.

### Website implementation

The VPTMdb web interface was written in the R programming language using the Rshiny web development framework [29]. The MySQL database management system was used to store structured PTM data. The base machine learning predictor (such as SVM) was supported by the caret R package [30]; the ROC curve was analysed using ROCR [31]; and MRMR and t-SNE were analysed using mRMRe [32] and Rtsne [33]. Software ggplot2 was used to plot beautiful pictures [34]. The website is free and can be browsed in most modern browsers.

## Acknowledgments

Thanks for the anonymous reviewers for their kind suggestions.

## Supporting information

**S1 Fig. Statistics of viral PTM data in VPTMdb**.

**S2 Fig. Viral protein kinase substrate motifs**. The HSV-1, and HSV-2 PTM amino acid residues were modified by US3 and UL13.

**S3 Fig. The results of KEGG and Gene Ontology enrichment analysis. S1 Table. The results of network analysis**.

**S2 Table. Training and independent datasets**.

**S1 Supporting Information. Supplementary materials**.

## Notes

http://vptmdb.com:8787/VPTMdb/

### Competing Interest Statement

The authors have declared no competing interest.

### Summary of Updates

Post-translational modifications (PTMs) plays an important role in the regulation of viral proteins; However, due to the limitation of data sets, there has been no detailed investigation of viral protein PTMs characteristics. In this manuscript, we collected experimentally verified viral protein post-translational modification sites and analysed viral PTMs data from a bioinformatics perspective. Besides, we constructed a novel feature-based machine learning model for predicting phosphorylation site. This is the first study to explore the roles of viral protein modification in virus infection using computational methods. The valuable viral protein PTM data resource will provide new insights into virus-host interaction.

