## Supplementary figures and images for "Bioinformatics analysis and collection of protein post-translational modification sites in human viruses"

### S1 Fig

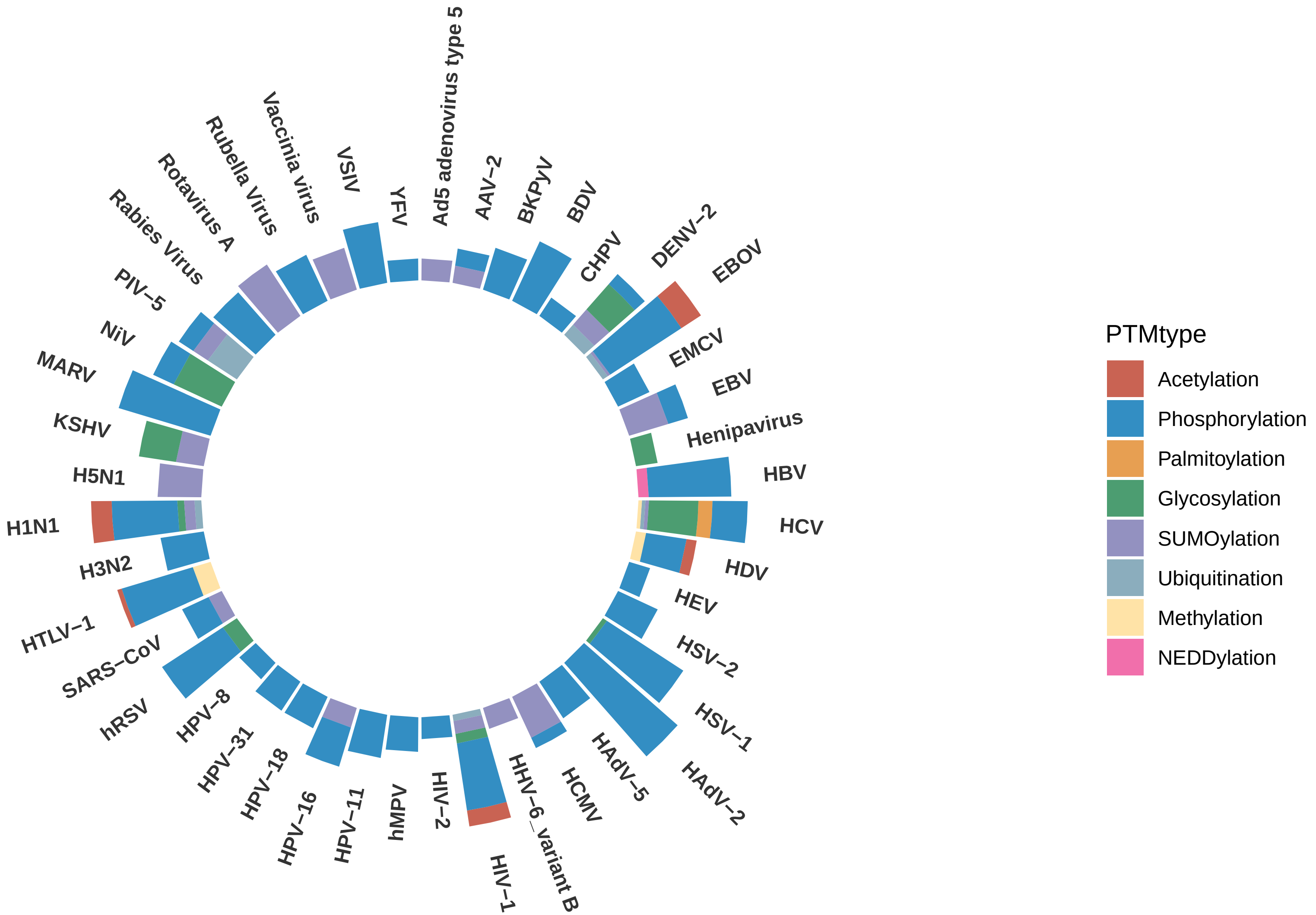

### S2 Fig

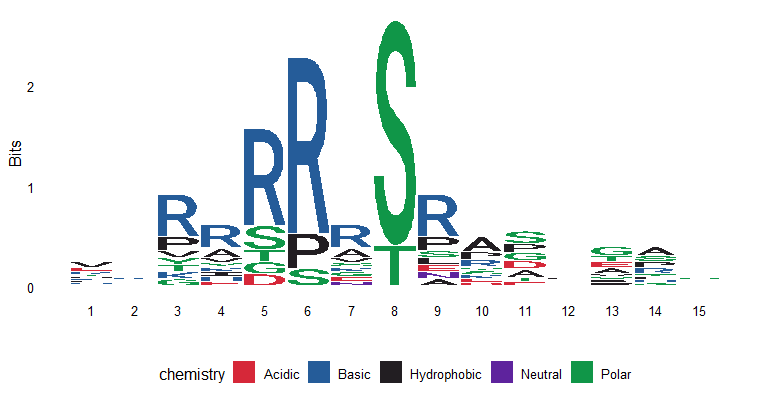

### S3 Fig

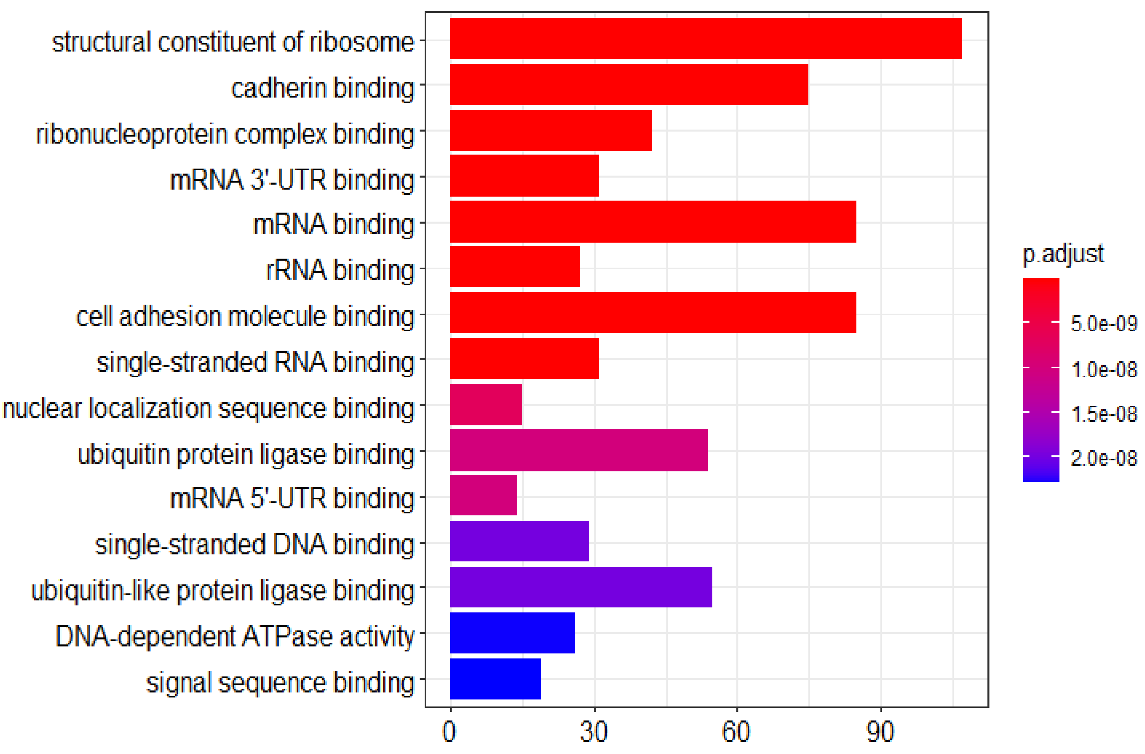

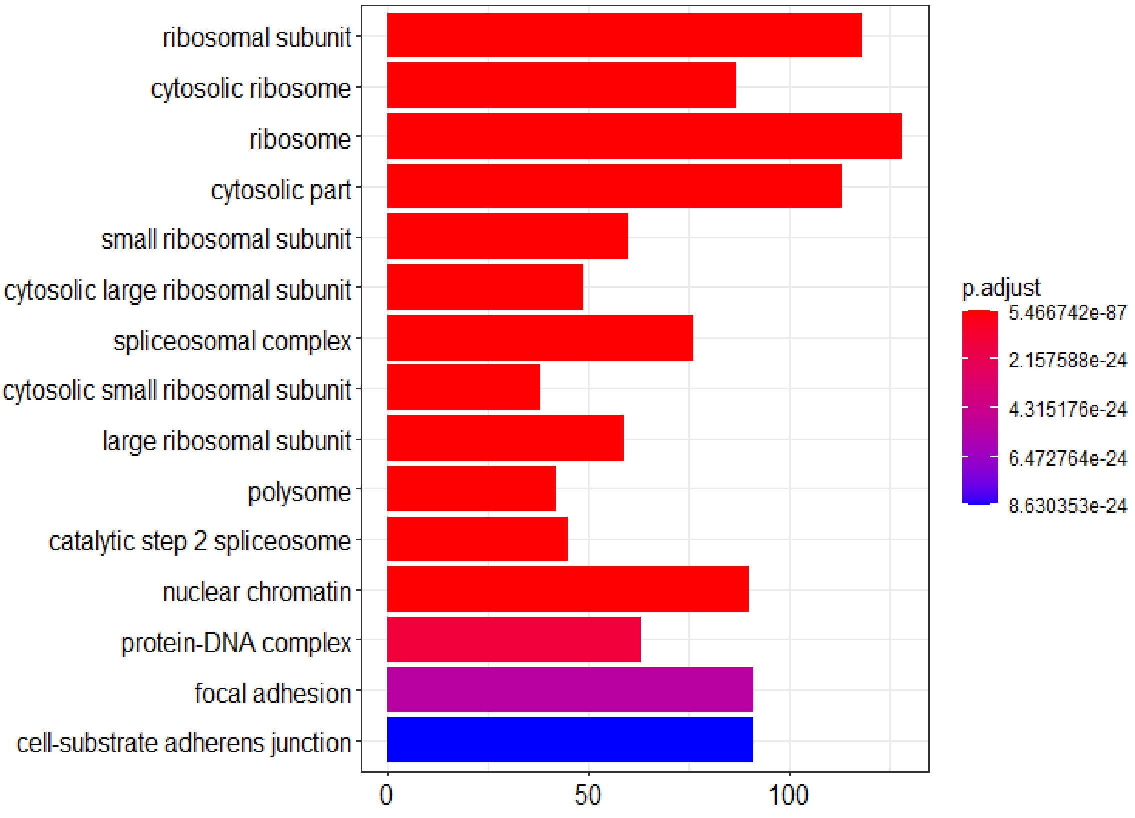

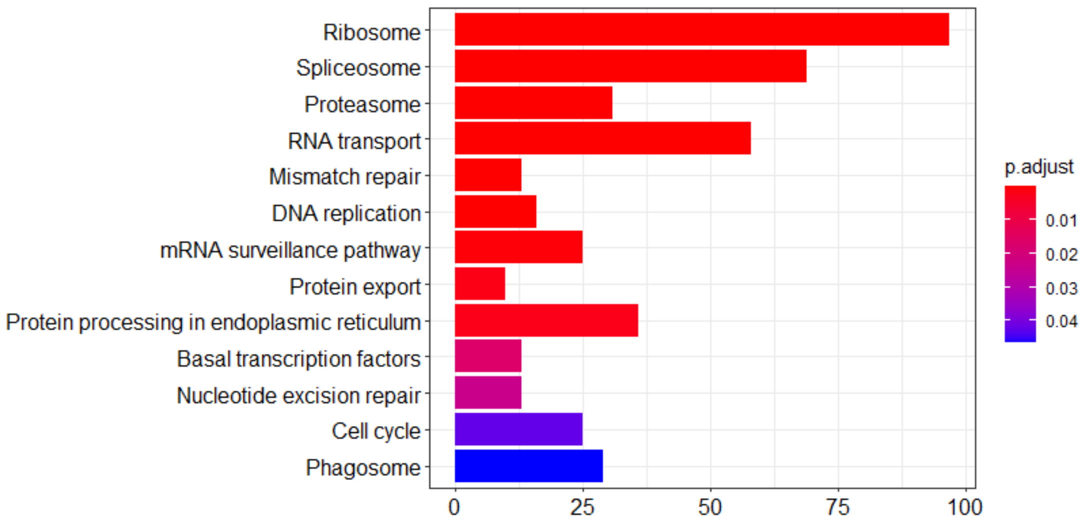
