## Supplementary material for "Bioinformatics analysis and collection of protein post-translational modification sites in human viruses": S1 Supporting Information

**Table of Contents**

**1. Supplementary Experimental Illustration**

Si1. Synthetic Minority Oversampling Technique (SMOTE)

Si2. Evaluation indicator

**2.** **Supplementary Table**

Table S1. The statistics of independent datasets in this study.

Table S2. The prediction performance of independent test between the original features and the combined features

Table S3. Different dimensions of combined features (AAC+ZSCALE), 5-fold cross-validation performance of three classifiers

Table S4. The independent test prediction performance of 68D.

**1. Supplementary Experimental Illustration**

**Si1. Synthetic Minority Oversampling Technique (SMOTE)**

SMOTE is an over sampling algorithm to construct balanced dataset developed by XXX.

The work flow is as followings:

(i) Defined the minority class samples as *T*_min_, *T*_min_ = {*X_1_, X_2_, X_3_, ... X_n_*}; Amount of SMOTE as N%; Number of nearest neighbors k.
(ii) For each *X_i_* in *T*_min_, computing the distance of it to all the samples in *T*_min_, sorting distance by near and far; Identified the k-nearest neighbors of each xi.
(iii) Selecting Xn, (n=1, ..., k) samples by from k-nearest neighbors according to the sampling ratio N%.
(iv) For each Xn, the new minority synthetic class sample can be defined as:

$$x\text{new}=xi+\text{rand}(\text{0-1})*\left| xi-xn \right|,i=1,\text{.}\text{.}\text{.},n$$

A simple example:

Consider a sample (6,4) and one of its k-nearest neighbor is (4,3),

$$x\text{new}=(6,4)+\text{rand}(0-1)*\left| (6,4)-(4,2) \right|$$

Where, rand (0-1) generates a random number between 0 and 1.

**Si2. Evaluation indicator**

We used five measures, i.e., sensitivity (Sn), specificity (Sp), F1-score and Matthews Correlation Coefficient (MCC) to evaluate the prediction performance in the present study.

Sensitivity or true positive rate (TPR) (percentage of correctly predicted phosphorylation sites):

.

Specificity (percentage of correctly predicted non- phosphorylation sites):

.

F1-score is a measure of a test's accuracy, the higher it is the more robust the classifier does

$$F1=\frac{2\text{TP}}{2\text{TP}+\text{FN}+\text{FP}}$$

MCC is a measure of the quality of binary classifications (Matthews, 1975), which is defined as

.

where TP, TN, FP*,* and FN represent the number of true positives, true negatives, false positives, and false negatives, respectively.

**2. Supplementary Table**

Table S1. The statistics of independent datasets in this study.

| **Independent subsets** | **positive** | **negative** |
| --- | --- | --- |
| Subsets1 | 52 | 53 |
| Subsets2 | 52 | 54 |
| Subsets3 | 52 | 54 |
| Subsets4 | 52 | 54 |
| Subsets5 | 52 | 53 |
| Subsets6 | 52 | 54 |
| Subsets7 | 52 | 54 |
| Subsets8 | 52 | 53 |
| Subsets9 | 52 | 53 |
| Subsets10 | 52 | 53 |

***Note***: Independent datasets for independent testing were constituted from 10 randomly selected negative subsets and 10 replicated positive datasets

Table S2. The prediction performance of independent test between the original features and the combined features.

| **Feature** | ***Sn*** | ***Sp*** | ***Acc*** | ***MCC*** | ***F1*** | ***AUC*** |
| --- | --- | --- | --- | --- | --- | --- |
| AAC | 0.769 | 0.548 | 0.658 | 0.324 | 0.689 | 0.7 |
| BINARY | 0.212 | 0.981 | 0.596 | 0.304 | 0.344 | 0.764 |
| ZSCALE | 0.231 | 0.979 | 0.605 | 0.319 | 0.369 | 0.715 |
| EGAAC | 0.635 | 0.712 | 0.673 | 0.348 | 0.657 | 0.72 |
| CTDD | 0.942 | 0.052 | 0.497 | -0.023 | 0.646 | 0.543 |
| CTDC | 0.788 | 0.54 | 0.664 | 0.339 | 0.698 | 0.722 |
| CTDT | 0.596 | 0.602 | 0.599 | 0.198 | 0.595 | 0.626 |
| CTriad | 0.385 | 0.873 | 0.629 | 0.297 | 0.508 | 0.726 |
| **ZSCALE+AAC** | **0.481** | **0.843** | **0.662** | **0.35** | **0.586** | **0.724** |
| ZSCALE+BINARY | 0.462 | 0.847 | 0.654 | 0.337 | 0.571 | 0.732 |
| ZSCALE+CTriad | 0.462 | 0.845 | 0.653 | 0.334 | 0.57 | 0.731 |
| ZSCALE+EGAAC | 0.442 | 0.845 | 0.644 | 0.317 | 0.553 | 0.727 |
| ZSCALE+CTDC+CTDT+CTDD | 0.5 | 0.744 | 0.622 | 0.252 | 0.567 | 0.706 |
| ZSCALE+EGAAC+BINARY+CTriad | 0.462 | 0.85 | 0.656 | 0.34 | 0.572 | 0.743 |
| ZSCALE+AAC+BINARY+Ctriad | 0.212 | 0.981 | 0.596 | 0.304 | 0.344 | 0.723 |

Table S3. Different dimensions of combined features (AAC+ZSCALE), 5-fold cross-validation performance of three classifiers.

| **Classifiers** | ***Dim*** | ***Sn*** | ***Sp*** | ***Acc*** | ***MCC*** | ***F1*** | ***AUC*** |
| --- | --- | --- | --- | --- | --- | --- | --- |
| **random forests** | 135 | 0.85 | 0.9 | 0.875 | 0.753 | 0.872 | 0.945 |
|  | 122 | 0.862 | 0.9 | 0.881 | 0.764 | 0.878 | 0.941 |
|  | 116 | 0.846 | 0.931 | 0.888 | 0.782 | 0.883 | 0.948 |
|  | 109 | 0.846 | 0.912 | 0.879 | 0.761 | 0.874 | 0.943 |
|  | 102 | 0.846 | 0.912 | 0.879 | 0.762 | 0.874 | 0.941 |
|  | 95 | 0.846 | 0.912 | 0.879 | 0.761 | 0.875 | 0.942 |
|  | 88 | 0.869 | 0.877 | 0.873 | 0.749 | 0.873 | 0.94 |
|  | 82 | 0.858 | 0.919 | 0.888 | 0.78 | 0.885 | 0.945 |
|  | 75 | 0.858 | 0.877 | 0.867 | 0.737 | 0.866 | 0.93 |
|  | 68 | 0.873 | 0.912 | 0.892 | 0.787 | 0.89 | 0.943 |
|  | 54 | 0.881 | 0.885 | 0.883 | 0.767 | 0.882 | 0.941 |
|  | 41 | 0.869 | 0.9 | 0.885 | 0.771 | 0.883 | 0.936 |
|  | 34 | 0.865 | 0.877 | 0.871 | 0.743 | 0.871 | 0.938 |
|  | 27 | 0.858 | 0.858 | 0.858 | 0.716 | 0.858 | 0.929 |
|  | 20 | 0.835 | 0.85 | 0.842 | 0.686 | 0.841 | 0.913 |
|  | 14 | 0.835 | 0.858 | 0.846 | 0.693 | 0.845 | 0.913 |
|  | 7 | 0.781 | 0.762 | 0.771 | 0.545 | 0.773 | 0.847 |
| **svm** | 135 | 0.904 | 0.873 | 0.888 | 0.78 | 0.891 | 0.958 |
|  | 122 | 0.904 | 0.862 | 0.883 | 0.77 | 0.887 | 0.954 |
|  | 116 | 0.904 | 0.862 | 0.883 | 0.77 | 0.887 | 0.954 |
|  | 109 | 0.908 | 0.869 | 0.888 | 0.782 | 0.892 | 0.953 |
|  | 102 | 0.892 | 0.865 | 0.879 | 0.762 | 0.881 | 0.949 |
|  | 95 | 0.885 | 0.85 | 0.867 | 0.738 | 0.87 | 0.948 |
|  | 88 | 0.9 | 0.842 | 0.871 | 0.746 | 0.875 | 0.948 |
|  | 82 | 0.904 | 0.846 | 0.875 | 0.753 | 0.879 | 0.955 |
|  | 75 | 0.915 | 0.823 | 0.869 | 0.744 | 0.875 | 0.948 |
|  | 68 | 0.904 | 0.846 | 0.875 | 0.754 | 0.879 | 0.946 |
|  | 54 | 0.9 | 0.796 | 0.848 | 0.703 | 0.855 | 0.939 |
|  | 41 | 0.896 | 0.788 | 0.842 | 0.691 | 0.851 | 0.928 |
|  | 34 | 0.892 | 0.8 | 0.846 | 0.696 | 0.853 | 0.921 |
|  | 27 | 0.885 | 0.788 | 0.837 | 0.679 | 0.844 | 0.905 |
|  | 20 | 0.854 | 0.765 | 0.81 | 0.624 | 0.817 | 0.881 |
|  | 14 | 0.796 | 0.723 | 0.76 | 0.522 | 0.768 | 0.843 |
|  | 7 | 0.696 | 0.646 | 0.671 | 0.344 | 0.679 | 0.753 |
| **naive bayes** | 135 | 0.788 | 0.431 | 0.61 | 0.227 | 0.671 | 0.746 |
|  | 122 | 0.769 | 0.638 | 0.704 | 0.414 | 0.725 | 0.804 |
|  | 116 | 0.742 | 0.727 | 0.735 | 0.473 | 0.737 | 0.836 |
|  | 109 | 0.762 | 0.758 | 0.76 | 0.523 | 0.759 | 0.845 |
|  | 102 | 0.765 | 0.754 | 0.76 | 0.523 | 0.76 | 0.844 |
|  | 95 | 0.762 | 0.754 | 0.758 | 0.52 | 0.758 | 0.845 |
|  | 88 | 0.769 | 0.769 | 0.769 | 0.543 | 0.769 | 0.849 |
|  | 82 | 0.762 | 0.769 | 0.765 | 0.534 | 0.764 | 0.851 |
|  | 75 | 0.773 | 0.765 | 0.769 | 0.541 | 0.77 | 0.853 |
|  | 68 | 0.758 | 0.773 | 0.765 | 0.532 | 0.763 | 0.853 |
|  | 54 | 0.792 | 0.769 | 0.781 | 0.563 | 0.784 | 0.854 |
|  | 41 | 0.792 | 0.792 | 0.792 | 0.585 | 0.793 | 0.859 |
|  | 34 | 0.785 | 0.788 | 0.787 | 0.574 | 0.787 | 0.86 |
|  | 27 | 0.796 | 0.769 | 0.783 | 0.566 | 0.786 | 0.855 |
|  | 20 | 0.788 | 0.781 | 0.785 | 0.57 | 0.786 | 0.845 |
|  | 14 | 0.746 | 0.758 | 0.752 | 0.504 | 0.751 | 0.826 |
|  | 7 | 0.708 | 0.665 | 0.687 | 0.374 | 0.694 | 0.773 |

Table S4. The independent test prediction performance of 68D.

| **Classifiers\evaluation** | ***sn*** | ***sp*** | ***Acc*** | ***MCC*** | ***F1*** | ***AUC*** |
| --- | --- | --- | --- | --- | --- | --- |
| random forests | 0.567 | **0.841** | **0.704** | **0.4272** | 0.656 | **0.744** |
| svm | 0.481 | 0.78 | 0.63 | 0.2754 | 0.564 | 0.705 |
| naïve bayes | 0.481 | 0.762 | 0.622 | 0.255 | 0.558 | 0.696 |
| PSI-blast | **0.717** | 0.65 | 0.676 | 0.358 | **0.681** | NA |
